# PTPN2-KO CAR-T Cells Demonstrate Enhanced Effector Function, CNS Infiltration, and Toxicity in a Non-Human Primate CAR-T Model

**DOI:** 10.1101/2025.09.17.676426

**Authors:** Francesca Alvarez Calderon, Ryan A Fleming, Marlana Winschel, Lev Gorfinkel, Rebecca Sutherland, Katherine Michaelis, James Kaminski, Elisa Rojas Palato, Kayleigh Ingersoll, Lorenzo Cagnin, Jennifer Lane, Victor Tkachev, Leslie S Kean, Ulrike Gerdemann

## Abstract

B-cell targeting CAR-T cell therapies achieve high remission rates, yet durable responses occur in fewer than 40% of patients. Deletion of negative T-cell regulators, such as PTPN2, a key inhibitor of TCR and cytokine signaling, represents a promising strategy to enhance the efficacy of CAR-T cells. While transfer of PTPN2 knockout (KO) T cells has demonstrated antitumor benefits in murine models, its impact on human-derived CAR-T cells and, importantly, the associated *in vivo* efficacy and toxicity remain unclear. Here, we demonstrate that PTPN2-KO human CD19 CAR-T cells exhibit enhanced cytokine production, cytotoxicity, TCR and CAR affinity and signaling, leading to superior *in vitro* elimination of leukemic cells with low CD19 expression. To assess *in vivo* efficacy and toxicity, we performed a dose-escalation study using a non-human primate (NHP) model of B-cell-targeting CD20 CAR-T cell therapy. We demonstrated that PTPN2-KO CD20 CAR-T cells exhibited superior *in vivo* expansion and B-cell depletion compared to WT CAR-T cells, in a dose-dependent manner. At the highest dose level, CAR-T expansion was associated with increased toxicities, particularly ICANS, compared to PTPN2 WT CD20 CAR-T cells driven by enhanced CNS-infiltration. Transcriptional profiling revealed a dominant effector and proliferative signature, with cytotoxic CNS-infiltrating CD8+ PTPN2-KO CAR-T cells implicated in ICANS pathogenesis. This study details the comprehensive evaluation of PTPN2-KO CAR-T cells in an immunocompetent model, demonstrating their enhanced on-target functionality, while highlighting increased toxicity risks, underscoring the need for rigorous preclinical assessment of potent genetic modifications in CAR-T therapy.

**Key points:**

- PTPN2-KO CAR-T cells exhibit enhanced effector function
- In a dose escalation study in rhesus macaques, PTPN2-KO mediated enhanced proliferation and CNS infiltration was associated with increased ICANS

## Introduction

Chimeric antigen receptor (CAR) T-cells have revolutionized the treatment of relapsed and refractory CD19+ leukemia and lymphoma, yet less than 40% of patients remain in long-term remission.^1,2^ Relapses have been linked to the upregulation of negative regulators of T-cell function,^2-4^ which dampens effective T-cell activation, limiting expansion and overall efficacy. Protein tyrosine phosphatases (PTPs) act as potent negative regulators of T-cell function by dephosphorylating signaling components downstream of the T-cell receptor (TCR) and by targeting JAK and STAT, thereby additionally attenuating cytokine signaling pathways^5,6^. The PTP non-receptor 2 (PTPN2) is highly expressed in rested and activated T-cells and other immune subsets including NK-cells, B-cells and myeloid cells^5^.

Evidence suggests that modification of PTPN2 may be a compelling approach to enhance CAR-T function. Monoallelic variants resulting in PTPN2 haploinsufficiency have been linked to early-onset autoimmune disorders, including inflammatory bowel disease, type I diabetes, rheumatoid arthritis, juvenile lupus, and Evans syndrome.^7^ In a murine tumor model of B16-OVA tumors, PTPN2-deleted OT-1 CD8 T-cells showed enhanced tumor invasion and increased activation marked by higher expression of CD25, Granzyme B, and IFN_γ_.^6^ Similarly, PTPN2-deleted mouse-derived HER2 CAR-T-cells improved tumor clearance of mammary tumors.^8^ Combined PTPN1/PTPN2 inhibition using a small molecule inhibitor improved T cell-mediated tumor clearance in murine models.^9^ These findings highlight the potential of PTPN2 inhibition to enhance tumor-directed T-cell function, providing a strong rationale for targeted genetic deletion in CAR-T cells. However, while PTPN2 emerges as a compelling candidate for clinical translation, rigorous preclinical evaluation to define both its therapeutic efficacy and the spectrum of potential toxicities is necessary.

CAR-T cell infusion is associated with two major toxicities: cytokine release syndrome (CRS) and immune-associated neurotoxicity syndrome (ICANS), which have limited the application of some of these therapies.^10^ Enhancing the efficacy of CAR-T cell therapies by targeting negative T-cell regulators has the potential to augment toxicities. Therefore, evaluating these potency-enhancing modifications in preclinical CAR-T cell models that accurately recapitulate human disease, reflect *in vivo* CAR-T cell efficacy, and enable the assessment of therapy-associated toxicities is a crucial safety measure.

Our lab previously described a CD20/B cell-directed CAR-T cell therapy model in rhesus macaques.^11,12^ This non-human primate (NHP) model assesses CAR-T cell efficacy by clearance of B cells, and reflects the clinical experience of early relapse in which patients show initial efficacy but limited CAR-T persistence (and reconstitution of B cells) as a harbinger of relapse.^2^ This feature enables the evaluation of increased CAR-T cell efficacy and key toxicities, including CRS and neurotoxicity, in an organism immunologically comparable to humans^11^.

In this study, we evaluated PTPN2-deleted B cell-targeting CAR-T cells both *in vitro* and in an *in vivo* NHP CAR-T model. Here we demonstrate that PTPN2-KO human CD19 CAR-T cells exhibit enhanced effector function and TCR affinity *in vitro*. Furthermore, in an *in vivo* dose-escalation study of CRISPR/Cas9-modified CAR-T cells in NHPs, PTPN2-KO CD20 CAR-T cells displayed increased proliferation and CNS-infiltration, albeit accompanied by dose-dependent high-grade CRS and ICANS, driven by a prominent proliferative effector T-cell transcriptional signature. These findings support PTPN2 editing as a strategy to enhance CAR-T potency in disease where CAR-T have been less effective.

## Methods

### Transduction and Expansion of NHP CD20 CAR-T cells

NHP CD20 CAR-T cells were generated as previously described.^11,12^ Viral transduction was performed 48-hours post-stimulation using spinoculation with a VSV-pseudotyped chimeric lentiviral/simian immunodeficiency virus encoding the CD20-CAR (containing 4-1BB and CD3*ζ*), and a truncated epidermal growth factor receptor (EGFRt) selection marker. After 24-hours, cells were electroporated using the Lonza 4D-Nucleofector System with a PTPN2-targeting RNA guide (5’-CCATGACTATCCTCATAGAG) or IDT control non-targeting guide Cas9 complex. After 4-6 days, stimulation beads were removed, and cells were enrichment using a PE-selection kit (StemCell Technologies). Final products were cryopreserved on day 10–14.

### Transduction and Expansion of human CD19 CAR-T cells

Human CD19 CAR-T cells were generated from healthy donor blood obtained under a Boston Children’s Hospital IRB-approved protocol. PBMCs were isolated and T-cells selected using the Pan T-cell negative isolation kit (Miltenyi Biotech). T-cells were activated with CD3/CD28-Dynabeads (Gibco) and transduced 48 hours later with a VSV-pseudotyped lentivirus encoding CD19 (FMC63) CAR with either 4-1BB or CD28 costimulatory domains and CD3ζ. Cells were electroporated with the same PTPN2-targeting or control gRNA used for NHP T-cells. Beads were removed after 4–6 days, and cells rested for 5–7 days before functional assays.

### Editing efficiency measurements in NHP and human edited T-cells

Genomic DNA was extracted from CAR-T cells using QuickExtract DNA extraction reagent (Lucigen Corp.) and amplified with PTPN2-specific primers (5’-GGCTGGGAAGATAAGTTTTGCTG, 5’-AAGGGGATGGAATGATATGAAAAGG). Sanger sequencing (Azenta/Genewiz) and ICE-Analysis (Synthego v3.0) were used to calculate the predicted KO score. Transcript expression was measured via SYBR Green qPCR with predesigned PTPN2 probes (IDT). Editing of the homologous PTPN1 locus was assessed by PCR amplification with PTPN1-specific primers (5’-GTCTGTTTCTTTGCTGGGTCTT, 5’-GGCGAAAGGAGTCTTGACTAAA) and analyzed via Sanger sequencing and ICE.

### Cytotoxicity and Cytokine Stimulation Assays

NALM6 cells were obtained from ATCC and grown per manufacturer’s instructions. CD19-low expressing NALM6^13^ cells were obtained from Dr. Robbie Majzner. Tumor cells were transduced to express the mcherry and luciferase under the control of EF1_α_ promoter

(Vectorbuilder), and flow sorted for purity. Tumor cells were co-cultured with CAR-T cells for 4-16 hours at the indicated effector-to-target ratios. Promega Bright-Glo luciferin solution was added at 1:1 volume, and luciferase expression analyzed on a Perkin-Elmer plate reader.

Cytokine stimulation assays were performed as previously described.^12^

### Jurkat Signaling Reporter Assays

Jurkat-Lucia NFAT and Jurkat-Dual NFκB/IRF reporter cells (Invivogen) were cultured per manufacturer instructions. These cells express Lucia luciferase under NFAT or NF-κB-responsive promoters, or SEAP under an ISG54-responsive promoter. Jurkat T-cells were edited as described to generate PTPN2-KO or control cells. CAR-Jurkat reporter cells were created via lentiviral transduction and flow sorting. T-cell activation was induced with anti-CD3 mAb, PHA, PMA/ionomycin, or NALM6 tumor cells overnight. Healthy donor CSF (Medix Biomedica) was used for assays as indicated. Transcription factor activity measured as secreted luciferase in supernatants was quantified using QUANTI-Luc 4 or Quanti-Blue reagents, subtracting background signal from media, and comparing fold-change to controls.

### NHP *In Vivo* T-cell Adoptive Transfer Studies

NHP experiments were approved by Massachusetts General Hospital and Biomere IACUC and conducted per NIH guidelines. Rhesus macaques received lymphodepleting chemotherapy with 40 mg/kg cyclophosphamide (Baxter) as previously described.^11^ NHP recipient characteristics are detailed in **Supplemental Table 1**. On day 0, CAR-T cells were thawed in RPMI media, and resuspended in PBS with 10% autologous serum for injection over 10-minutes. CAR-T cell doses ranged from 1×10^4^–6×10^6^/kg viable cells.

Recipients received prophylactic ceftazidime and vancomycin (maintaining 5–10 mcg/mL vancomycin levels) and, during lymphopenia (ALC <1000 cells/mL), weekly cidofovir with probenecid and acyclovir until recovery. Keppra (15 mg/kg) was administered from day 0 until CRS/ICANS resolution. Neurologic monitoring (activity level, movement/speed, presence of tremors, gait abnormalities, and seizures) mirrored patient evaluations.^14^ **(Supplemental Table 2)**. CRS was treated with up to two doses of tocilizumab (∼10 mg/kg rounded to nearest vial size), and severe ICANS was managed with dexamethasone (1–4 mg/kg) and diazepam (0.5 mg/kg) as indicated.

### Brain Dissociation and T-cell Purification

NHP brains were harvested at necropsy following perfusion with 3–5 L PBS until organ pallor and clear fluid were observed. White matter was sectioned under veterinary supervision. Brain tissues were dissociated in PBS with 150 KU DNase (Sigma-Aldrich) and 1 mg/mL type-I collagenase (Invitrogen) at 37 °C for 1 hour. The digested material was filtered through metal and then nylon strainers (100–40 µm). Leukocytes were isolated using a double-layer Percoll gradient (GE Healthcare) and used for flow cytometry, single-cell sequencing, or cryopreserved in 10% DMSO/90% FBS.

### Sample Preparation for Single Cell Transcriptomic Analysis

Cryopreserved NHP products and post-infusion PBMCs, brain, and CSF cells were thawed or prepared fresh, stained with Live/Dead stain (Invitrogen), CD4, CD8, CD3, and EGFR-t antibodies, and sorted via flow cytometry for single-cell sequencing. Samples were processed using the Next GEM Single Cell 5’ v2 Sample Index Plate TT (10x Genomics). Details of scRNA-Seq analysis methods are in the Supplemental Methods.

## Results

### PTPN2-KO CAR-T cells exhibit enhanced *in vitro* effector function

We first identified a CRISPR/Cas9 guide that would selectively and effectively delete PTPN2 in primary human T-cells in a region with low homology to PTPN1 (**Supplemental Fig.1A-B**), achieving a mean editing efficiency of 73.5+/-2.9% in primary T and CAR-T cells measured by interference analysis using Synthego ICE deconvolution (**Fig.1A, Supplemental Fig.1A)** and transcriptional analysis **(Supplemental Fig.1B)**.

**Figure 1.**
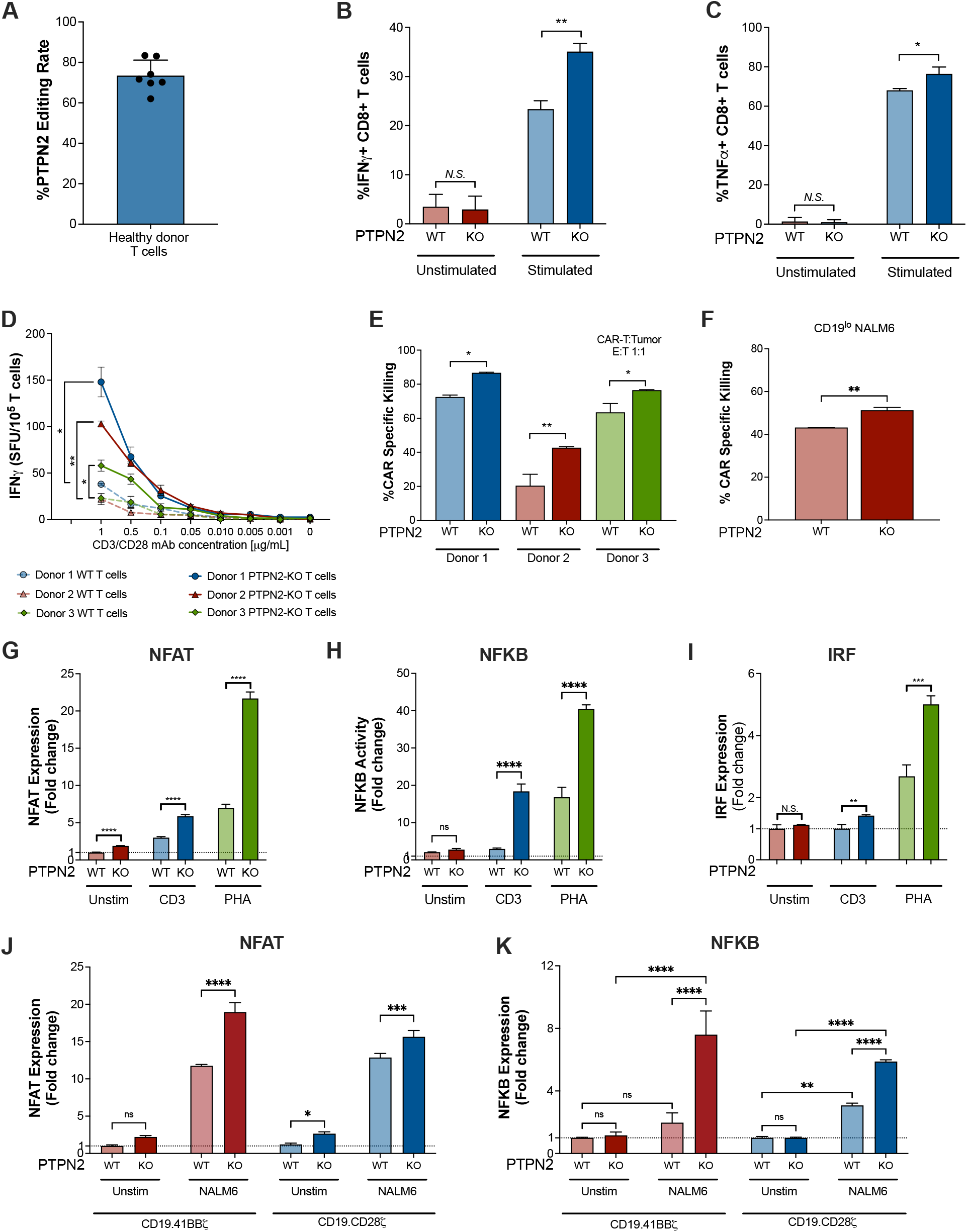
Human PTPN2-KO CAR-T cells require a lower activation threshold and exhibit enhanced killing and activation. **A**. CRISPR/Cas9 PTPN2 gene editing efficiency in healthy human T cells quantified by Synthego ICE deconvolution from Sanger sequencing spectra. Each dot represents a different T cell donor. **B-C**. PTPN2 WT or KO T cells were stimulated with PHA/Ionomycin and stained for IFN_ψ_ (B) and TNF_α_ (C). Fraction of positive cells of total CD8+ T cells is shown. **D**. IFN_ψ_ secretion in human PTPN2 WT and KO T cells measured by ELISpot immunoassay following limiting dilution stimulation with CD3 agonistic monoclonal antibody for 18-hours. **E**. CAR-T cell cytotoxicity against NALM6 tumor cells 4 hours after co-culture with PTPN2-WT or KO CD19.28ζ CAR-T cells at an effector-to-target ratio of 1:1. **F**. CAR-T cell cytotoxicity against CD19^low^ NALM6^15^ tumor cells 4 hours after co-culture with PTPN2-WT or KO CD19.28ζ CAR-T cells at an effector-to-target ratio of 1:1. **G-I**. Fold change in NFAT (G), NFĸB (H), and IRF (I) activity in PTPN2-WT or KO Jurkat reporter cell lines after activation with CD3 monoclonal antibody or PHA for 16h. **J-K**. Fold change in NFAT (J) and NFĸB (K) in PTPN2-WT or KO CD19.41BBζ (red) and CD19.CD28ζ (blue) CAR Jurkat reporter cell lines after CAR-specific stimulation with NALM6 for 18 hours. Representative results from at least 2 independent experiments with different healthy T cell donors as indicated. Unpaired t-test, * indicates p<0.05, ** p<0.01 and *** p<0.001.

To assess the impact of PTPN2 deletion on human T-cell effector function, we assessed cytokine production and cytotoxic activity against leukemic cells using *in vitro* functional assays. When compared to CD8 T-cells edited with a non-targeting control guide that retained expression of the endogenous wild type PTPN2 (PTPN2-WT), PTPN2-knockout (PTPN2-KO) CD8 T-cells exhibited significantly increased IFN-γ production (35.07% vs 23.37%, SE 1.38, p=0.001) and elevated TNF-α levels (76.47% vs 68.07%, SE 2.06, p= 0.15) **(Fig.1B,C)** upon stimulation with PMA and ionomycin. PTPN2 deletion disrupts the dephosphorylation of critical downstream molecules in the TCR signaling pathways, potentially augmenting TCR avidity. To evaluate this, PTPN2-KO and PTPN2-WT T-cells were stimulated with limiting dilution concentration of a CD3 agonistic antibody. PTPN2-KO cells demonstrated significantly higher IFN-γ production at lower stimulus concentration **(Fig.1D, Supplemental Fig.1D)**, consistent with a significantly reduced activation threshold for PTPN2-KO cells due to higher TCR signaling strength.

To assess the cytotoxic potential of PTPN2-KO T-cells, we generated human-derived CD19-41BB-CD3ζ PTPN2-KO and PTPN2-WT CAR-T cells and co-cultured them with a CD19-expressing B-ALL cell line (NALM6). PTPN2-KO CD19 CAR-T cells demonstrated enhanced *in vitro* target cell lysis compared to PTPN2-WT CAR-T cells (1.2-2-fold higher killing in KO vs WT depending on the donor, **Fig.1E)**. Similarly, PTPN2 deletion was associated increased CAR functional avidity, as demonstrated by the superior killing of CD19 low-expressing NALM6 cells^15^ by PTPN2-KO CAR-T cells relative to their WT counterparts (43.2 vs. 51.3%, p=0.003) **(Fig.1F, Supplemental Fig.1E)**. These findings highlight not only the enhanced effector and cytotoxic capabilities of PTPN2-KO CAR-T cells but also their increased TCR and CAR avidity, resulting in superior killing of leukemia cells with low antigen density.

### PTPN2-KO directly regulates NFAT, NF-κB and IRF downstream programs in CAR-T cells

To determine whether PTPN2 deletion broadly enhances downstream signaling from multiple T-cell activation pathways, including NFAT, NF-κB, and interferon regulatory factors (IRFs), we generated a panel of PTPN2-KO and PTPN2-WT (edited with control non-targeting guide RNAs) Jurkat reporter cell lines. In the absence of stimulation, PTPN2-KO Jurkat NFAT-luciferase cells exhibited a modest yet significant increase in endogenous NFAT activity relative to PTPN2-WT Jurkat cells **(Fig.1G)**. Upon TCR stimulation using either CD3-agonistic antibody or phytohemagglutinin (PHA), a selective T-cell mitogen, this difference was further amplified, leading to substantial upregulation of NFAT, NF-κB, and IRF activity in PTPN2-KO reporter cell lines **(Fig.1G-I)**.

To assess whether PTPN2-mediated regulation of these signaling pathways would also modulate CAR activation, and to assess the impact of different costimulatory domains, we transduced PTPN2-KO Jurkat reporter cells with either a CD19-41BB-CD3ζ or CD19-CD28-CD3ζ CAR. Upon CAR stimulation using NALM6 cells, we observed enhanced NFAT and NF-κB activity independent of the costimulatory domain (**Fig.1H-I)**. These results indicate that PTPN2-deletion leads to broad and sustained enhancement of both TCR- and CAR-driven signaling.

### Single cell transcriptomic analysis reveals a strong proliferative effector signature of PTPN2-KO CAR-T cell products

To evaluate the efficacy and toxicity of PTPN2-KO CAR-T cells in a clinically and immunologically translatable model, we conducted a dose-escalation study of PTPN2-KO CD20/B cell-directed CAR-T cells in our previously established NHP CAR-T model.^11,12^ PTPN2 is highly conserved between humans and rhesus macaques, with 98.96% sequence homology in coding regions (Supplemental Fig. 4), enabling the use of the same exon 2-targeting CRISPR/Cas9 guide for both species (Fig. 2A), and ensuring a translatable preclinical assessment **(Fig.2A)**. Underscoring the role of PTPN2 in CAR-T cell activation *in vitro* and *in vivo*, 10x single-cell RNA sequencing (scRNAseq) analyses of historical WT CAR-T NHP recipients demonstrate that PTPN2 is highly expressed on both CAR+ and CAR^Neg^ CD8 and CD4 T-cells in NHPs^12^, and upregulated during *in vitro* manufacturing and *in vivo* expansion (**Fig.2B**).

**Figure 2.**
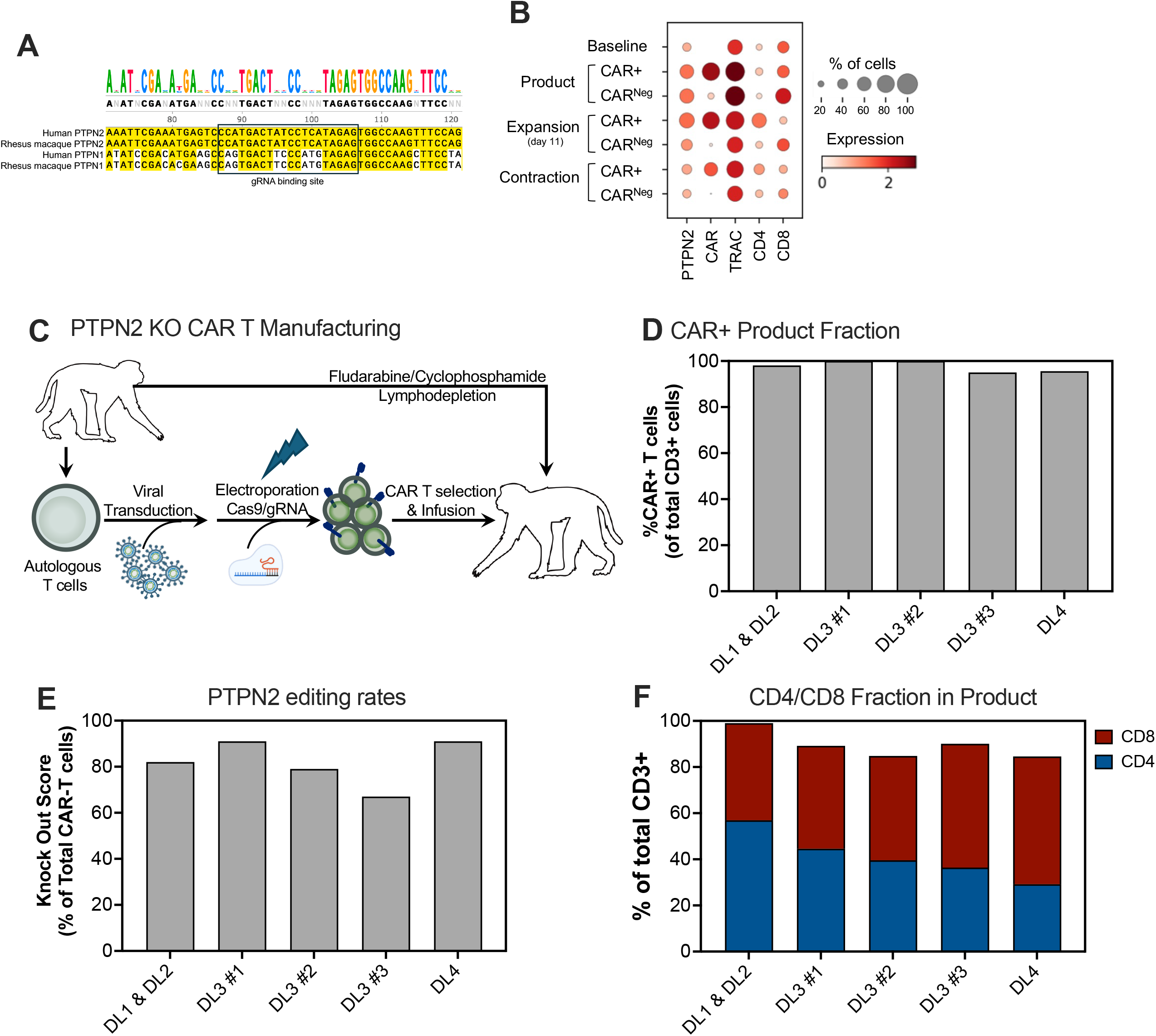
Rhesus macaque PTPN2-KO CAR-T cells manufacturing process. **A**. Gene alignment of PTPN1 and PTPN2 in human and rhesus macaque at the PTPN2 gRNA binding site. **B**. *PTPN2* expression in single cell transcriptomic analysis of PTPN2-WT CAR-T cell recipients compared to canonical T cell markers (*TRAC, CD4 and CD8*). Bubble size represents fraction of cells, and color intensity expression level. **C**. Manufacturing process of autologous CD20-directed BBζ PTPN2-KO CAR-T cell, and subsequent infusion into lymphodepleted recipients. **D**. Fraction of CAR+ T-cells measured by EGFRt expression in PTPN2-KO CAR-T manufactured products. **E**. PTPN2 editing rates in PTPN2-KO CAR-T infusion products quantified by Synthego ICE deconvolution from Sanger sequencing spectra. **F**. CD4 and CD8 fraction of total CD3+ T cells in manufactured products.

To generate PTPN2-KO CAR-T cells, T cells were transduced with CD20-BBζ CAR co-expressing EGFRt and nucleofected with PTPN2 gRNA/Cas9 RNP. To avoid infusion of non-CAR-expressing PTPN2-KO T-cells, which could induce autoimmunity as seen in murine PTPN2-KO T-cell transfer experiments^16,17^, CAR-T cells were selected by magnetic beads prior to cryopreservation **(Fig.2C)** ensuring a CAR fraction of 95.6-100% **(Fig.2D)** in 5 products generated. CAR-T had a KO score of 67-91% **(Fig.2E)**, and varying CD4/CD8 ratios **(Fig.2F)**.

To elucidate transcriptional differences in the generated PTPN2 vs WT CAR-T cell products, we performed 10x single-cell RNA sequencing (scRNAseq) analysis on PTPN2-KO CAR-T cells (n=3) and PTPN2-WT CAR-T cells (n=2) after *in vitro* manufacturing **(Fig.3A)**. After alignment and quality control, we recovered 27,120 CAR-T cells. Unsupervised clustering **(Fig.3B-F)** yielded a total of 8 clusters, annotated based on expression of the top 50 enriched transcripts, including CD8-clusters (proliferating, cytotoxic and early memory clusters), CD4-clusters (memory/naïve) and mixed CD8/CD4-clusters (memory/naïve) **(Fig.3F)**. CAR-T cells were subsetted by CD8 and CD4 expression for pseudo-bulk analysis comparing KO vs WT CAR-T cells. A small subset of cells expressed both CD8 and CD4 but with limited numbers, no further analysis was performed on this population. Scored gene lists from differentially expressed transcripts in CD8+ PTPN2-KO vs WT-CAR-T cells were analyzed using ClusterProfiler for the C2 and C7 compartments of the Human MSigDB Collections Gene-Set Enrichment Analysis (GSEA). CD8+ PTPN2-KO CAR-T cells demonstrated enrichment of multiple cell cycle pathway entries, with normalized enrichment score (NES) ranging 1.319-1.657 (p-adjusted<0.05), a TCR pathway with an NES 1.753, p-adjusted 0.04, and activation/memory pathways, with NES ranging 1.783-1964, p-adjusted<0.005, with enrichment towards a more naïve/memory phenotype characterized by expression of *CCR7, SIT1, TXNIP, ZFP36L2, SPRY1, GRB10* **(Fig.3F-I)**. Additional enriched pathways are shown in **Supplemental Fig.4**. Taken together, these data are consistent with PTPN2-KO CAR-T cells exhibiting higher proliferation and TCR activation signatures while retaining a more naïve/memory phenotype similar that previously described with the use of PTPN1/2 small molecule inhibitors.^9^

**Figure 3.**
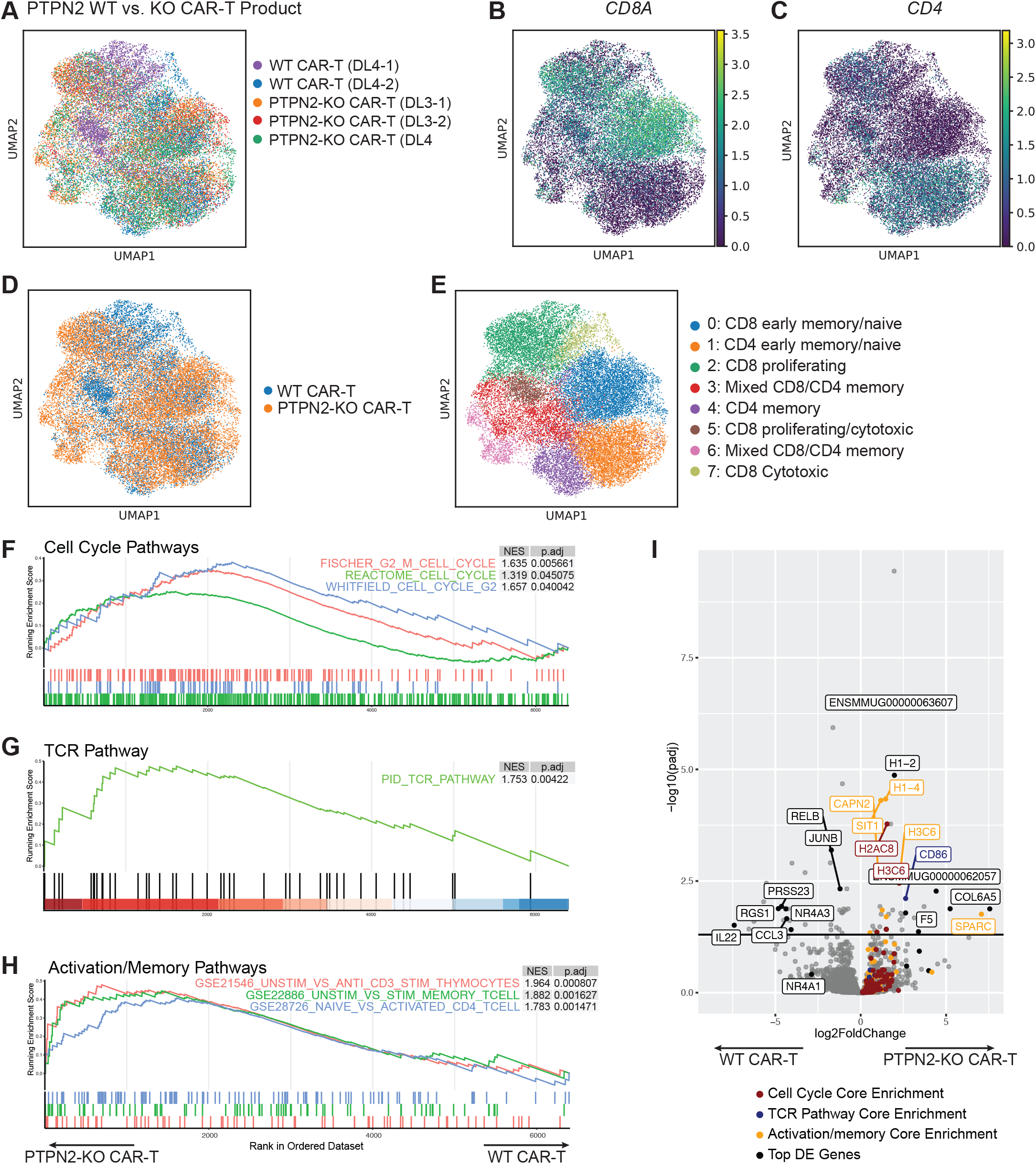
Single cell transcriptomic analysis of infusion products reveals PTPN2-KO CAR-T cells have enhanced proliferation and T cell activation. **A**. UMAP embedding of product CAR-T cells after *in vitro* manufacturing and selection for CAR expressing T cells by bead selection of EGFRt+ cells. **B-C**. UMAP embedding showing normalized expression of *CD8A* (B), and *CD4* (C). **D**. UMAP embedding showing PTPN2 WT vs KO CAR-T products. **E**. UMAP embedding showing cluster annotations based on CD4/CD8A expression, and evaluation of top 50 highly enriched genes in each cluster. **F-H**. Gene Set Enrichment Analysis of pseudobulk CD8+ CAR-T cells compared to cell cycle pathways (F), TCR activation pathways (G) and activation/memory pathways. Normalized enrichment scores and adjusted p-values are denoted next to reference gene set. **I**. Volcano plot showing differential expression of transcripts. Cell cycle, TCR pathway, and activation/memory core enrichment genes are colored as indicated.

### PTPN2-KO CAR-T cells exhibit enhanced *in vivo* proliferation in the NHP CAR-T model

PTPN2-KO CAR-T cell doses were infused in a dose escalation study, ranging from 6×10^4^ CAR-T cells/kg (dose-level 1, DL1) to 6×10^6^ CAR-T cells/kg (DL4) **Supplemental Table 1)**. PTPN2-WT control animals received DL4 (n=2). NHP recipients received cyclophosphamide lymphodepletion prior to intravenous administration of freshly thawed CAR-T cell product^11^ **(Fig.2C)**.

The clinical outcomes of the dose escalation study are summarized in **Fig.4A**. PTPN2-KO CAR-T cells at DL1 (1×10^4^ CAR-T cells/kg, n=1) showed no measurable CAR-T expansion, B cell aplasia, or clinical/laboratory findings associated with CAR-T cell expansion. DL2 (1×10^5^ CAR-T cells/kg, n=1) demonstrated 4.9% PTPN2-KO CAR-T cells on day 7 but no associated B cell aplasia or clinical/laboratory signs of CRS or ICANS **(Fig.4A)**. The same animal received DL1 and DL2 41 days apart, following repeated lymphodepletion, which may have limited the expansion. At DL3 (3×10^6^ CAR-T cells/kg, n=3), all animals demonstrated CAR-T expansion (max expansion 18.6±6.8% CAR-T, **Fig.4B**) and B cell aplasia lasting a mean of 32.7±9 days (ranging from day 27 to 43, **Fig.4C)**. All animals developed laboratory signs of CRS, including elevation in CRP, Ferritin, and LDH **(Fig.4D-F)**; and clinical signs of CRS including fever, edema, and tachypnea; and ICANS, including tremors, unstable gait, and decreased responsiveness (**Fig.4A, Supplemental Table 1**). One animal in PTPN2-KO DL3 developed severe CRS with respiratory distress, and ICANS with clinical symptoms of encephalopathy **(Supplemental Table 1)**; and required treatment with Tocilizumab and Dexamethasone **(Fig.4A)**. This coincided with peak CAR-T cell expansion of 27.4% CAR+ on day 8.

**Figure 4.**
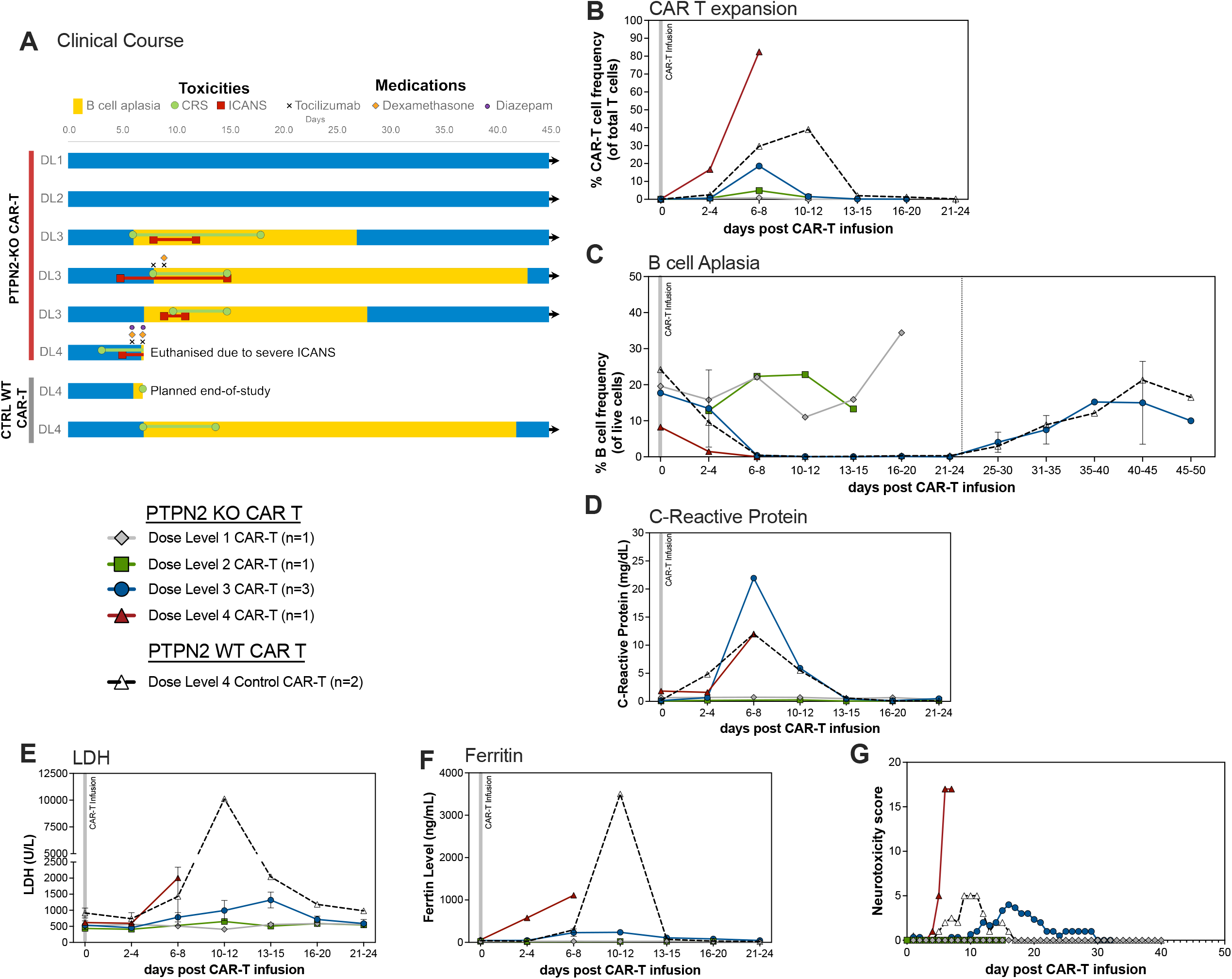
PTPN2-KO CAR-T cells demonstrate increased *in vivo* expansion and toxicities. **E**. Swimmer plot summarizing dose escalation *in vivo* study in rhesus macaques. B cell aplasia is denoted by yellow bar, clinical course including CRS and ICANS by lines, and symbols indicate medication administration, See Table 1 for additional details. **B**. CAR-T expansion in blood measured by flow cytometry. CAR+ fraction was determined by expression of EGFRt of total CD3+ T cells. **C**. B cell aplasia defined as <1% of PBMCs. **D-G**. Laboratory evaluation of CRS including levels of C-reactive protein (D), LDH (E) and Ferritin (F). **G**. ICANS grading for CAR-T recipients.

Administration of dexamethasone led to rapid contraction of circulating CAR-T cells and resolution of ICANS and CRS.

A single NHP received PTPN2-KO CAR-T cells at DL4 (6×10^6^ CAR-T cells/kg). This animal exhibited rapid CAR-T expansion (16.7% CAR+ T-cells on day 5) and higher CAR-T expansion (82.4% of total T-cells were CAR+ on day 7), which was more expansion than PTPN2-WT DL4 doses, which exhibited maximum expansion of 42.3+/-3.3% CAR+ cells on day 7-9 (**Fig.4B**).

The DL4 PTPN2-KO CAR-T experiment also demonstrated earlier onset of B cell aplasia (day 3 vs 5-7) (**Fig.4C**). This animal exhibited severe CRS and ICANS unresponsive to treatment with Tocilizumab and repeated doses of Dexamethasone, necessitating humane euthanasia **(Fig.4A)**. Due to dose-limiting toxicities observed at DL4, no additional animals were infused with PTPN2-KO cells at DL4 or higher. Importantly, NHPs in our prior work^11^ receiving control CAR-T at higher cell doses up to 1.2×10^7^ CAR-T/kg did not experience severe CRS or ICANS symptoms (**Supplemental Table 1, Fig.4B**), suggesting that increased potency of PTPN2-KO CAR-Ts was associated with increased toxicity. Of note, all PTPN2-KO CAR-T recipients (DL2-4) with evidence for CAR-T expansion demonstrated a predominance of CD8 PTPN2-KO CAR-T cells when compared to the corresponding infusion products (**Supplemental Fig.3C**).

### CNS-infiltrating PTPN2-KO CAR-T cells are predominantly cytotoxic CD8 CAR-T cells

NHP recipients of PTPN2-KO CAR-T cells exhibited high-grade ICANS, particularly the DL4 recipient **(Fig.4G, Supplemental Table 1)**. Clinical and pathological analysis revealed widespread cerebral edema and CAR-T cell infiltration into the brain parenchyma. CAR-T cell fractions in CNS compartments (white matter, brainstem, periventricular matter, cerebellum, and CSF) were significantly higher in PTPN2-KO recipients compared to PTPN2-WT recipients (CNS-to-blood ratio: 1.47 vs. 0.26, **Fig.5A)**. Additionally, since the infusion product for the PTPN2-KO recipients had a small fraction of PTPN-WT CAR-T, we evaluated the PTPN2-KO enrichment by measuring the fraction of edited cells in sorted cells from blood and brain-infiltrating CAR-T cells. Compared to the infusion product (63% PTPN2-KO CAR-T cells), we observed a progressive increase in the proportion of these cells in the blood (72%) and brain parenchyma (84%), suggesting preferential expansion and CNS-infiltration of PTPN2-KO CAR-T cells (**Fig.5B**).

**Figure 5.**
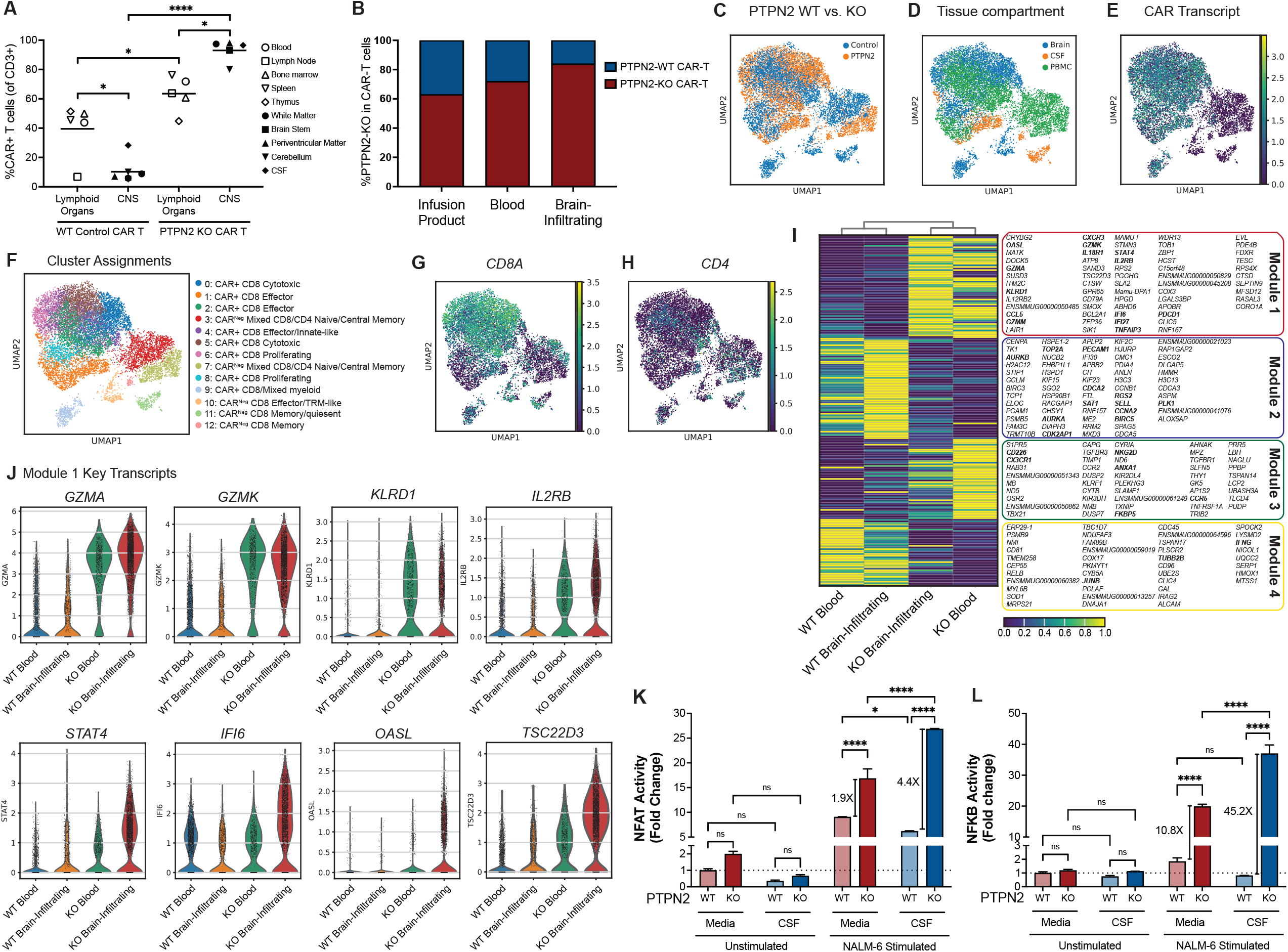
PTPN2-KO CAR-T cells show enhanced infiltration into CNS. **A**. Flow cytometry analysis of CAR+ cells of total CD3+ T cells in peripheral lymphoid organs and central nervous system at time of maximum expansion in recipients receiving DL4 PTPN2 WT vs. KO CAR-T cells. Historical control WT CAR-T cell recipient published^11^ included for reference. One-way ANOVA, followed by 2-way student t-tests. **B**. PTPN2 editing rates in fluorescently cell sorted CAR+ CD3+ T cells at time of product thaw, and maximum expansion in PTPN2-KO DL4 CAR-T cell recipient. Editing rates calculated by Synthego ICE deconvolution from Sanger sequencing spectra. **C-D**. UMAP embedding of T cells defined by expression of *CD3E* or *TRAC* transcripts colored by recipient (C) and tissue type (D). **E**. Cluster cell type annotations based on CD8A/CD4 expression, CAR Transcript expression, and top 50 highly enriched genes in each cluster. **G-H**. UMAP embedding showing normalized expression of *CD8A* (G) and *CD4* (H) transcripts. **I**. Heatmap showing normalized expression of differentially expressed transcripts in PTPN2 WT vs KO blood and brain samples, grouped by unsupervised hierarchical clustering into 4 distinct modules. Gene transcripts are listed in order. **J**. Violin plots show expression of transcripts in Module 1 upregulated in CD8+ PTPN2-KO brain and blood. **K-L**. NFAT and NF-ĸB Activity in PTPN2 WT and KO CAR-Jurkat reporter cells upon stimulation with NALM6 in media vs. healthy donor CSF. Representative results from 2 independent experiments. Unpaired t-test, * indicates p<0.05, ** p<0.01 and *** p<0.001.

To assess the transcriptional profile of brain-infiltrating PTPN2-KO CAR-T cells, we conducted scRNA sequencing on paired blood, CSF, and CNS-infiltrating CAR-T cells from the DL4 PTPN2-KO CAR-T cell recipients and compared it to a PTPN2-WT CAR-T cell recipient, which exhibited low-grade ICANS at the time of necropsy. T cells were identified by expression of *CD3E* or *TRAC*, and subsequently clustered using Scanpy PCA and UMAP analysis. We recovered 12,734 cells, including 9,819 expressing the CAR transcript **(Fig.5E)**. Unsupervised clustering of T cells identified 12 clusters including 5 CAR^neg^ CD8 and mixed CD8/CD4 clusters, 7 CAR+ CD8+ clusters, and 1 CAR+ mixed CD8/CD4 cluster **(Fig.5F)**. scRNA-Seq analysis was consistent with the flow cytometric findings that the brain-infiltrating PTPN2-KO CAR-T cells (cluster 0,2,4,5,6) were predominantly CAR+ CD8+ T cells **(Fig.5G)**. We performed single-cell differential expression analysis of PTPN2-KO vs PTPN2-WT cells from the blood and brain (the two compartments with sufficient CAR+ T cells numbers for analysis) and identified 214 significant differentially expressed transcripts among the comparisons **(Fig.5I)**. Hierarchical clustering demonstrates a pattern of 4 distinct sets of transcripts predominantly driven by PTPN2 editing, denoted in the heatmap as Module 1-4 **(Fig.5I)**. Module 1 comprised transcripts upregulated in PTPN2-KO brain-infiltrating CAR-T and to a lesser extent in PTPN2-KO blood CAR-T cells, including cytotoxic (*GZMA, GZMK, KLRD1*), cytokine signaling (*IL2RB, STAT4*,

*IL12RB2, IL18R1, CCL5*) and interferon response genes (*IFI6, IFI27* and *OASL*) (**Fig.5J and Supplemental Fig.5)**. Module 2 comprised transcripts upregulated in PTPN2-WT brain-infiltrating CAR-T cells and included cell cycle (*AURKA, PLK1, CENPA*) with a more naïve phenotype (*SELL*) (**Supplemental Fig.5)**. Module 3 included transcripts upregulated in PTPN2-KO blood CAR-T cells and was dominated by tissue trafficking effector markers (*CX3CR1, CCR5, CCR2, TXNIP,TBX21*) and co-stimulation/activation receptors (*NKG2D, CD226, SLAMF1*). Module 4 comprised transcripts upregulated in PTPN2-WT blood CAR-T and to a lesser extent in brain-infiltrating WT CAR-T including *IFNG*, and activation/transcription factor signaling (*RELB, JUNB*), without other canonical effector or activation markers (**Supplemental Fig.5)**. In the PTPN2-KO groups, we identified several steroid inducible transcripts (*TSC22D3, ANXA1, FKBP5*) consistent with administration of dexamethasone in this recipient. Importantly, despite steroid administration, the PTPN2-KO CAR-T cells retained effector and cytotoxic markers. Taken together, PTPN2-KO CAR-T cells exhibit enhanced cytotoxicity, cytokine signaling, trafficking and co-stimulation/activation.

We hypothesized that PTPN2-KO CAR-T cells would have enhanced activation in the CNS microenvironment and thus enrich in the CNS. To evaluate the functional effect of CSF on PTPN2-KO CAR-T cell activity, we stimulated PTPN2-KO and WT CAR-Jurkat cells in either cell culture media or healthy donor human CSF, and measured NFAT and NF-κB activity, two key transcription factors downstream of CAR activation. When cultured in CSF, WT CAR-T cells show a decrease in NFAT activity and modest non-significant decreased in NF-κB activity upon activation with tumor cells. In contrast, when cultured in CSF, PTPN2-KO CAR-Jurkat cells show higher NFAT and NF-κB activity after stimulation (Difference in NFAT activity 4.4X after stimulation in CSF compared to 1.9X in media, and NF-κB activity 45.2X compared to 10.8X, **Fig.5K-L**), suggesting that PTPN2 deletion may be able to overcome a postulated more suppressive CNS microenvironment^18^.

## Discussion

In this study, we have detailed the immunologic and clinical phenotype of PTPN2-KO CAR-T cells. These studies demonstrate that PTPN2 deletion can enhance the *in vitro* and *in vivo* effector function of B-cell targeting CAR-T cells and enhance CNS-infiltration of CAR-T cells. However, *in vivo*, the increased effector function and CNS-infiltration was also associated with exacerbated toxicity, particularly ICANS, underscoring the potential tradeoffs that may occur with enhanced CAR-T function. PTPN2 serves as a critical negative regulator of both TCR and cytokine signaling pathways, making its deletion a promising strategy to enhance CAR-T cell efficacy. Previous studies have demonstrated that PTPN2 knockout can augment antitumor responses in murine models^6^. However, preclinical evaluations of PTPN2-KO CAR-T cells have primarily focused on efficacy, lacking a comprehensive assessment of their toxicity profile, which has been challenging to model in murine studies. Most CAR-T animal studies have relied on transplanted human tumor xenografts into NSG mice^19^, but the limited ability to evaluate toxicity in these models, and the lack of long-term follow-up due to CAR-T rejection and development of graft-vs-host disease from xeno-transplantation have limited the clinical evaluation of T cell-enhancing modifications.^20^ While syngeneic models provide a more physiologically relevant microenvironment, fundamental differences between murine and human CAR-T cells present challenges for direct clinical translation. Notably, none of these models fully recapitulate the spectrum of CAR-T cell-associated toxicities observed in patients. Our previously established NHP model of B-cell-targeting CAR-T cell therapy is able to accurately recapitulate both therapeutic efficacy and associated toxicities in a highly homologous animal model. We have now demonstrated that PTPN2-KO CAR-T cells exhibit enhanced expansion, effector function and CNS-infiltration but were also associated with an increased incidence and severity of CRS and ICANS.

In experiments using PTPN2-WT CAR-T cells in NHP, toxicities were restricted to elevated inflammatory markers, indicative of CRS, accompanied by mild, self-limited neurological symptoms, including tremors that did not necessitate supplemental therapeutic intervention^11^. In contrast, administration of PTPN2-KO CAR-T cells, even at lower doses compared to previous studies with WT CAR-Ts^11^, led to high-grade ICANS, accompanied by enhanced CAR-T cell infiltration into the CNS. Our single-cell transcriptomic analyses demonstrate that CNS-infiltrating CD8^+^ PTPN2-KO CAR-T cells adopt an enhanced cytotoxic transcriptional program, potentially contributing to the increased incidence of ICANS. This PTPN2-specific effect is further supported by *in vitro* findings, where PTPN2-KO CAR-T cells display enhanced activation when cultured in CSF compared to standard media. Further investigation using the NHP model may provide critical insights into the pathogenesis of ICANS.

This study highlights a potential distinction between systemic inhibition of PTPN1/2 and targeted PTPN2 deletion in antigen-specific effector cells. Systemic administration of PTPN1/2 inhibitors is hypothesized to carry a risk of autoimmune disease, given PTPN1/2 expression across multiple immune cell populations, and autoimmunity reported in patients with PTPN2 haploinsufficiency^7^ and murine models^16,17^. In contrast, in our study, targeted PTPN2 deletion was restricted to CAR-T cells, and did not induce autoimmune-related clinical issues, suggesting that CAR-T restricted PTPN2 editing may offer a more precise therapeutic approach compared to systemic inhibition.

While the enhanced CNS-infiltration of PTPN2-KO CAR-T likely contributed to the heightened ICANS, and may constrain the clinical utility of CAR-Ts directed against some B cell malignancies, their enhanced effector function could be advantageous in settings where CAR-T cell efficacy remains suboptimal, such as with low antigen-expressing tumors, and CAR-T cells for solid tumors. Furthermore, the preferential CNS-infiltration and CNS effector function of PTPN2-KO CAR-T cells may offer a therapeutic benefit for primary or secondary CNS malignancies, where enhanced CNS homing and local CAR-T effector function are desirable, and currently addressed by repeated dosing of intracerebroventricular CAR-T cells.^21,22^ However, the observed dose-dependent toxicity underscores the necessity for rigorous dose-escalation studies to ensure safety in PTPN2-KO CAR-T cell therapies.

In conclusion, this study demonstrates the potential of genetic modifications to enhance CAR-T cell function while emphasizing the importance of using clinically relevant animal models to assess the translational viability of CAR-T cell engineering strategies.

## Data sharing statement

scRNA-sequencing data will be available through the Gene Expression Omnibus (GEO) database.

## Acknowledgments

We would like to thank the veterinary staff in the Center for Comparative Medicine at Massachusetts General Hospital for care and supportive services provides to our primate recipients. We would like to thank members of the Cellular Therapeutics and Systems Immunology Lab, and our colleagues Natalie Collins, Jared Rowe, and Marc Schwartz for their thoughtful scientific discussions and support of this project. This work was supported by National Institutes of Health R01-HL095791 (LSK), P01-HL158504 (LSK), U19-AI051731 (LSK), NIH K12-HD052896 (FAC), NIH T32-HL007574 (FAC, UG, LG, RS), ASTCT New Investigator

Award (VT), NMDP Amy Strelzer Manasevit Research Grant (VT).

## Authorship Contributions

FAC, UG, and LSK designed experiments. FAC, UG, RF, MW, LG, EJRP, KAM performed research. FAC, UG, RF, MW analyzed data. FAC, UG, and LSK wrote the manuscript. All authors critically reviewed and approved the content of the manuscript.

