## Supplemental Materials for "PTPN2-KO CAR-T Cells Demonstrate Enhanced Effector Function, CNS Infiltration, and Toxicity in a Non-Human Primate CAR-T Model"

### 10X Single-cell RNA sequencing analysis pipeline

#### Alignment of scRNA-seq Samples

GEX libraries were aligned using the 'Cellranger multi' pipeline (v7.0.0) with 32 CPU's and 128 GB of RAM. Samples were aggregated with the 'Cellranger aggr' pipeline (v7.0.0) with arguments '--normalize=none --nosecondary'. These were aligned against a custom reference transcriptome created with the Cellranger makeref command using rhesus macaque genome assembly Mmul_10 as the reference genome and Ensembl v114 as the reference set of genomic annotations for the NHP samples representing the most up to date reference genome for rhesus macaque at time of analysis. In addition to the gene set from Ensembl, sequences for the two CAR transcripts used in the study were added for the custom NHP transcriptome.

#### Initial QC and Filtering

Fastqc was used to assess the quality of the sequencing, and the output metrics from Cellranger were used to assess the quality of the libraries. The samples were processed through CellBender for removal of ambient RNA and empty droplets then aggregated into an annData object and loaded into Scanpy. Data from across all batches were integrated using batch correction by sequencing batch. We performed QC using selected key metrics and removed samples with <10% of transcripts mapped to mtDNA along with those that had > 350 genes detected. We then divided the dataset into the Product and non-Product for separate analyses as previous analysis has shown when clustered together the largest driving force in difference is whether a cell is product or non-product. We retained highly variable genes (HVGs) by concatenating the top 4,000 HVGs from the Product samples and the top 7500 HVGs from the non-product samples, resulting in 8708 unique HVGs.

#### Identification of Latent Space, Clusters, and Normalized Counts with scVI

Scvi-tools was used to fit a variational autoencoder (VAE) on raw transcript counts separately on each of two (product and non-product) objects using the parameters n_layers=2, n_latent=30, gene_likelihood="nb", batch_key="Seq_Batch". The "get_latent_representation" function from scVI was used to get the latent space from the VAE for each dataset then used Scanpy to run the neighbors, UMAP and Leiden functions using default settings for initial clustering and dimensionality reduction. The "get_normalized_expression" function from scVI with library size set to 10,000 was used to obtain normalized counts for the cells.

**Identification of Product Signature with scVI**

The product samples were clustered using scVI as described above. The data was subsetted into CD4 only and CD8 only by expression of CD4, CD8A and CD8B for all downstream analysis steps. A small subset of cells expressed both CD4 and CD8 as has previously been described in rhesus macaques, these cells were not carried to downstream analysis due to low cell count. Pseudobulk analysis of PTPN2-KO vs Control PTPN2-WT was executed by summing raw gene counts within each animal and using DESeq2 in R for the CD4 only and CD8 only compartments. Genes from the PTPN2-KO CD8 compartment were analyzed for GSEA using the ClusterProfiler R package. Ranking for this GSEA analysis was performed by generating a score for each gene by dividing their lfc_median by the lfc_std (error) to ensure robust results. These scored gene lists were then used with ClusterProfiler for the C2 and C7 compartments of the Human MSigDB Collections. Major pathways identified including TCR pathway, Cell Cycle pathways and Naïve/memory pathways are shown. Differentially expressed transcripts from the Pseudobulk analysis are also represented by a volcano plot with this gene list using ggplot visualization of the lfc_mean and -log10(probab_not_de) with genes from relevant GSEA pathways highlighted by color and selected significant genes, including the top 10 significant DE genes by lfc in either direction as well as those relevant to significant GSEA pathways, are labeled.

**PTPN2-KO and PTPN2-WT DL4 Recipients Clustering with scVI**

The clustered annData object containing all biological samples was filtered to T cells using expression of CD3E and/or TRAC. Initial clustering was examined and clusters with low representation of CD3E and TRAC were dropped to preserve only T cells for analysis. The object was then re-clustered and separated into CD4 and CD8 compartments as described above for the product object. Clustering showed a clear delineation between clusters with relatively high CAR transcript expression (>45%) and those without (<10%). Cells contained in clusters with high CAR transcript expression were further examined for single cell differential expression analysis. Single cell differential expression analysis was executed on several biologically relevant compartments by utilizing only specific samples as idx1 and idx2 as arguments in the vae.differential_expression function using the VAE from initial biological sample clustering. The CD4 and CD8 positive cells of each of the blood and brain compartments were separately compared between the two animals against one another to elucidate transcriptional differences between PTPN2-WT and KO. A matrixplot of relative gene expression level (normalized by gene) is shown highlighting the significant, highly differentially expressed genes between the different compartments with significance determined by > 1 or <-1 lfc_median, bayes_factor>3, is_de_fdr_0.05 == True, and nonzero proportion >.1. UMAPs of normalized gene expression of selected differentially expressed gene genes are shown.

### Supplemental Figures

**Supplemental Figure 1**. Human PTPN2-KO CAR-T cells show selective editing at PTPN2 and require a lower activation threshold.

**Supplemental Figure 2.** Needleman-Wunsch global alignment demonstrate high homology between human and NHP PTPN2 coding regions.

**Supplemental Figure 3.** Rhesus macaque PTPN2-KO CAR-T cells demonstrate enhanced expansion *in vivo*.

**Supplemental Figure 4.** Gene Set Enrichment Analysis of pseudobulk CD8+ CAR-T cells

**Supplemental Figure 5.** PTPN2-KO CAR-T cells show district transcriptomic signatures in CNS compared to blood.

### Supplemental Tables

**Supplemental Table 1.** Summary of NHP recipients reported in this study

**Supplemental Table 2.** Neurotoxicity score / ICANS for NHP recipients

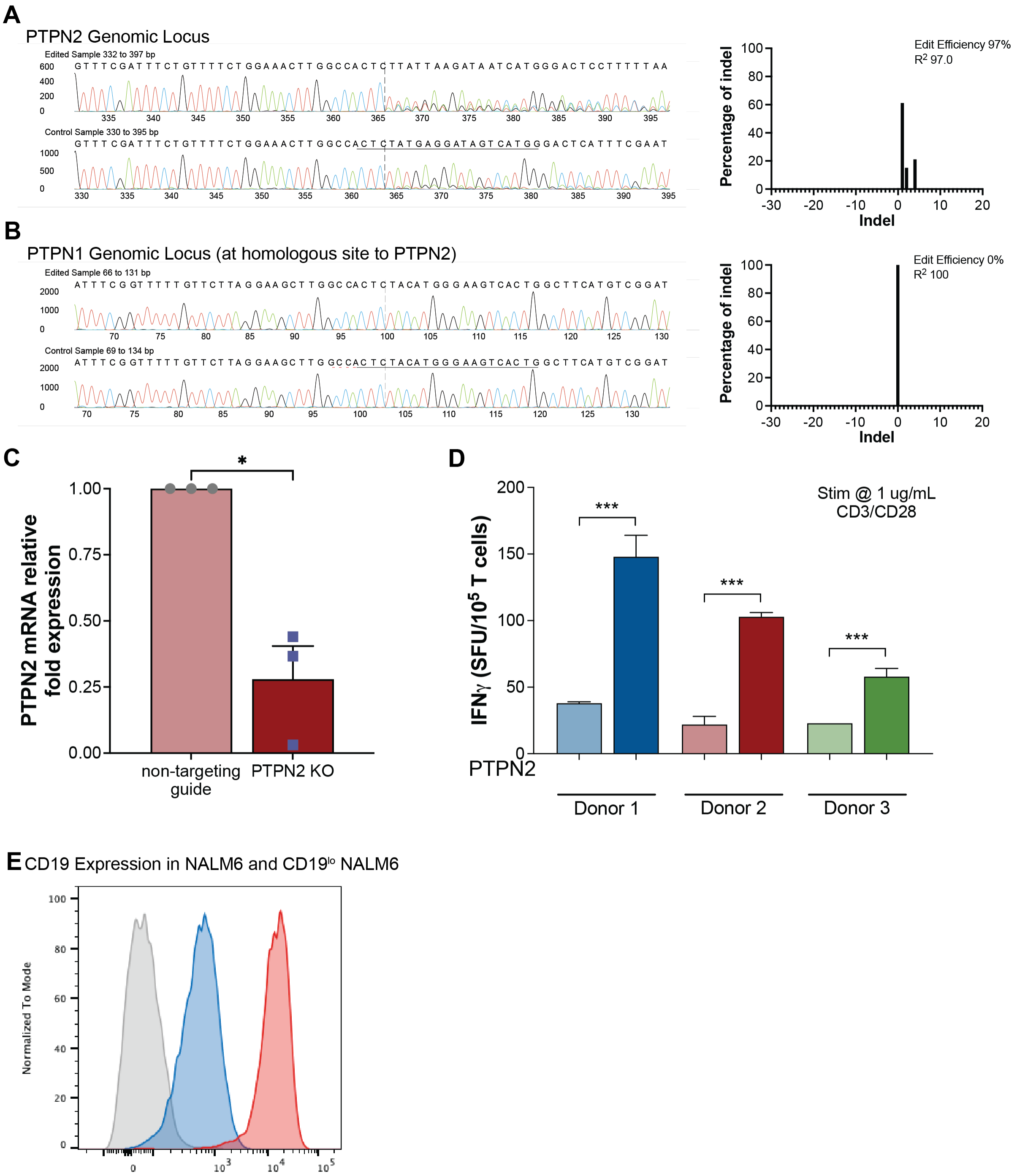


**Supplemental Figure 1**. **Human PTPN2-KO CAR-T cells show selective editing at PTPN2 and require a lower activation threshold. A-B.** Representative Synthego ICE deconvolution of Sanger sequencing spectra to determine PTPN2 (A) and PTPN1 (B) editing rates. Shown is PTPN1 homologous loci as PTPN2 gRNA binding site. **C.** PTPN2 relative mRNA expression normalized to GAPDH in PTPN2-KO vs. WT human T cells. Each dot represents a unique healthy donor. **D.** IFNγ secretion in human PTPN2 WT and KO T cells measured by ELISpot immunoassay after stimulation with 1 μg/mL of CD3 monoclonal antibody for 48-hours. **E.** CD19 expression in CD19$^{low}$ NALM6 cell line compared to WT NALM6 tumor cells.

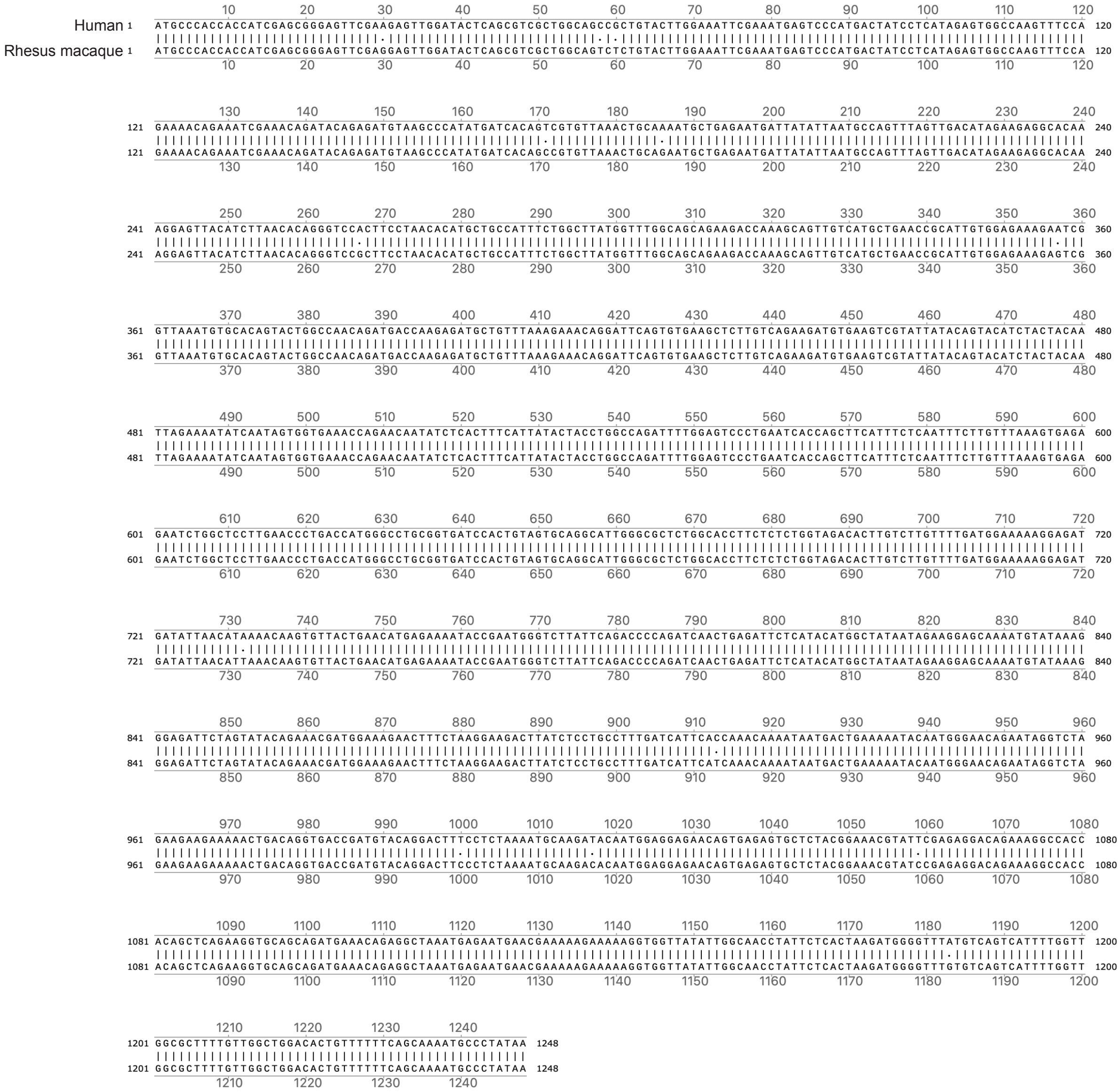


**Supplemental Figure 2. Needleman-Wunsch global alignment demonstrate high homology between human and NHP PTPN2 coding regions.** Human PTPN2 coding region sequence is shown in the top sequence while rhesus macaque sequence in bottom sequence. Dots indicate difference in sequences leading to 3 amino acid substitutions at p.P20S, p.P305S and pS334P.

### Supplemental Figure 3

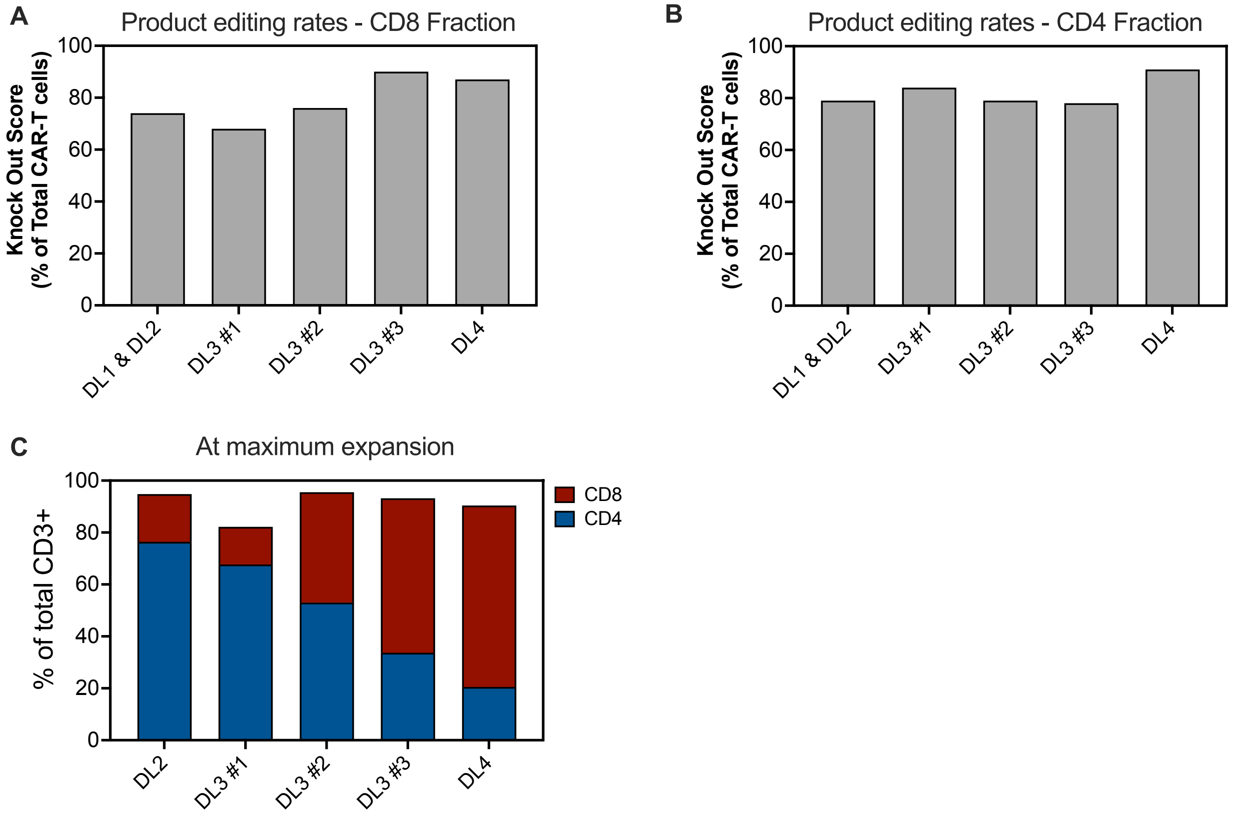


**Supplemental Figure 3. Rhesus macaque PTPN2-KO CAR-T cells demonstrate enhanced expansion *in vivo*. A-B.** PTPN2 editing rates in PTPN2-KO CAR-T infusion products quantified by Synthego ICE deconvolution from Sanger sequencing spectra in CD8 (A) and CD4 (B) T cells. **C-D.** Fraction of CD4 and CD8 CAR-T cells in infusion products (C) and at the time of maximum *in vivo* expansion (D). Cells are gated on CAR+ CD3+ lymphocytes.

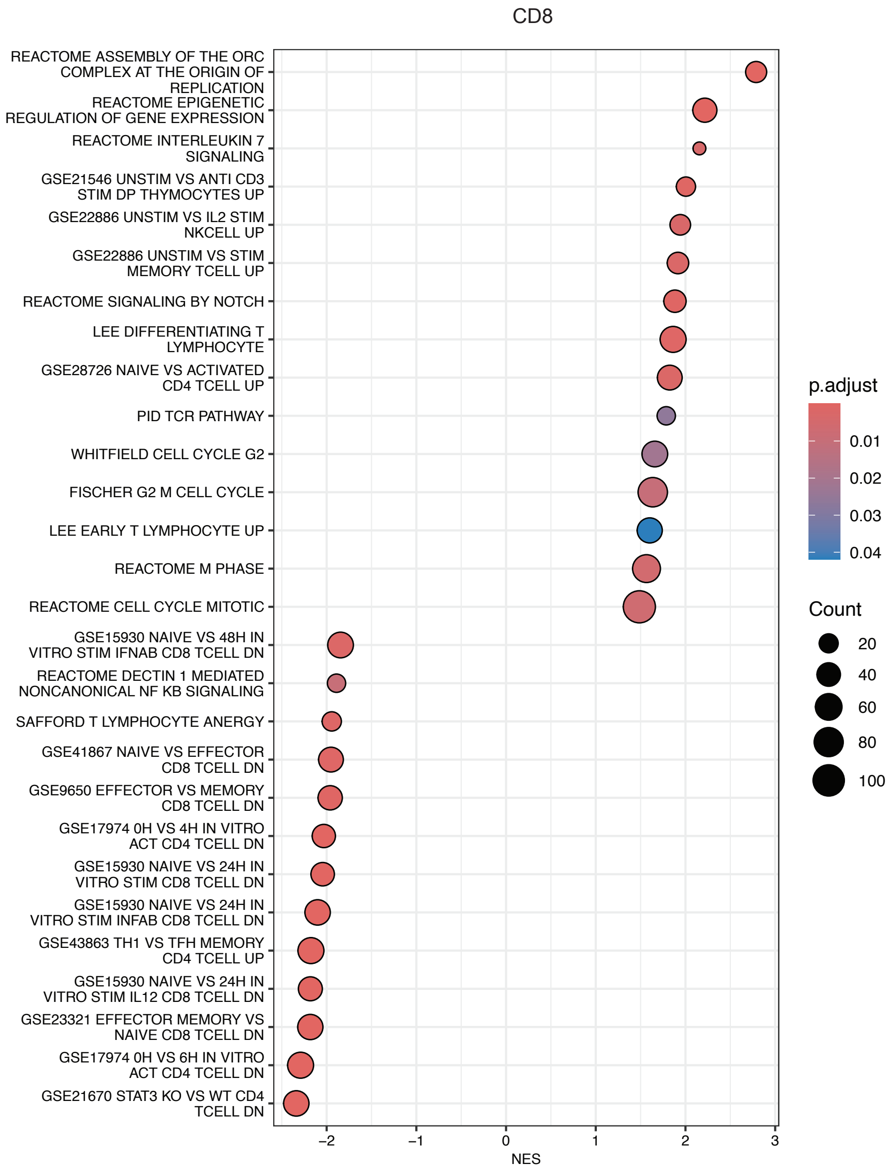


**Supplemental Figure 4. Gene Set Enrichment Analysis of pseudobulk CD8+ CAR-T cells**. selected key T cell pathways are shown. Normalized enrichment scores of key pathways are shown in Figure 3F-H.

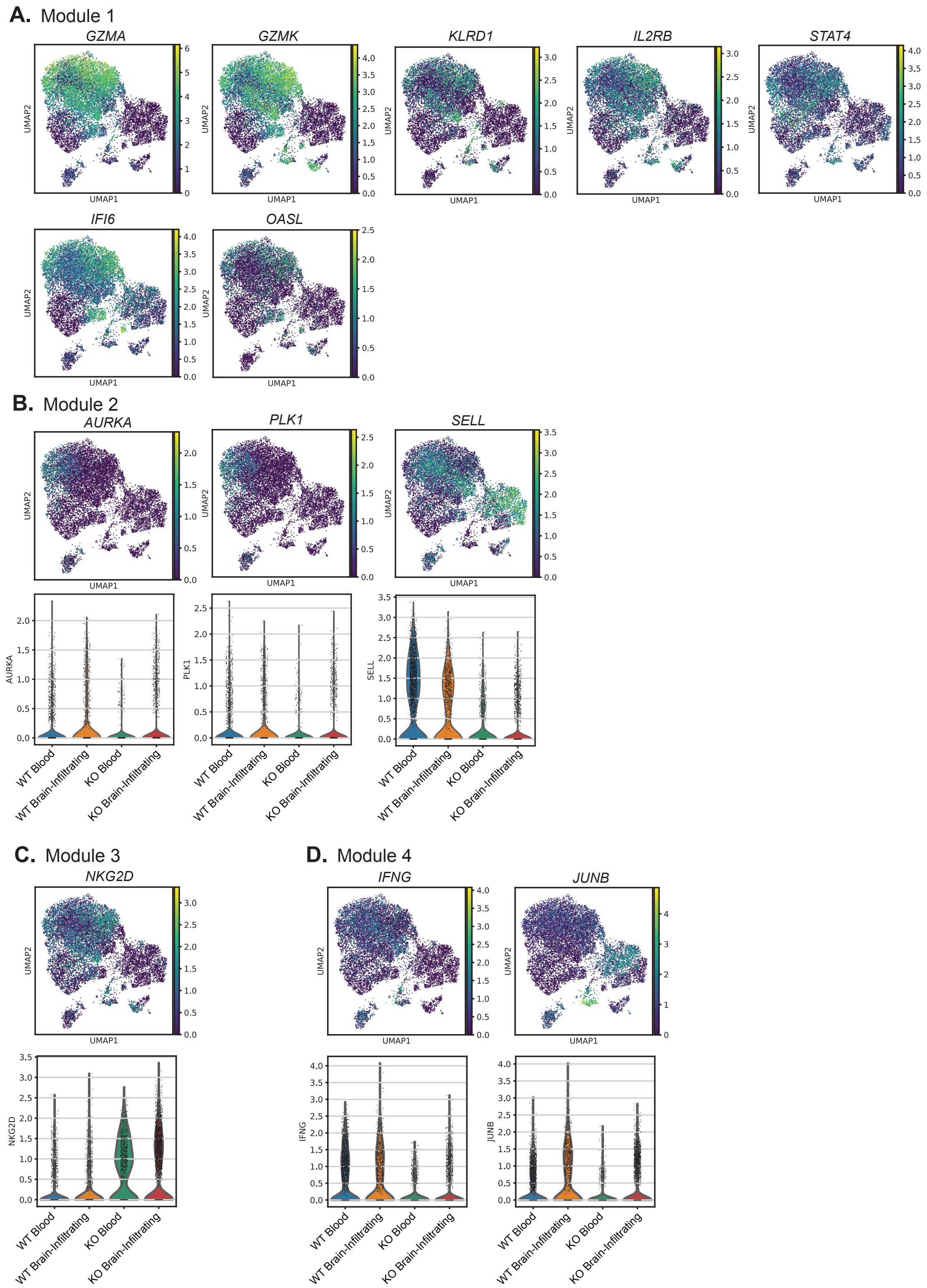


**Supplemental Figure 5. PTPN2-KO CAR-T cells show district transcriptomic signatures in CNS compared to blood. A.** UMAP embedding of key transcripts identified in Module 1 (related to Figure 5I). **B-D.** Violin plots and UMAP embedding of transcripts identified in Module 2 (B), Module 3 (C) and Module 4 (D).

**Supplemental Table 1.** Summary of NHP recipients reported in this study

| Animal ID | Age | Sex | Weight | CAR-T cell dose/kg | Max CAR-T cell expansion | Max Grade CRS | Max Grade NHP ICANS[1] | Treatment | Endpoint day | Reason for endpoint |
|---|---|---|---|---|---|---|---|---|---|---|
| CTRL WT CAR-T DL4 | 1y 5m | Male | 7.9 kg | 6.00E+06 | 45.60% | 1 | 3 | Keppra prophylaxis | d7 | Planned study endpoint |
| CTRL WT CAR-T DL5 | 5y 3m | Female | 7 kg | 1.23E+07 | 39% | 1 | 5 | Keppra prophylaxis | d60 | |
| PTPN2-KO CAR-T DL1, DL2 | 4y 3m | Male | 7.2 kg | 1.00E+04, 1.00E+05 | 0.8%, 4.9% | 0 | 0 | Keppra prophylaxis | d60 post DL2 | |
| PTPN2-KO CAR-T DL3#1 | 4y 11m | Male | 8.6 kg | 1.00E+06 | 23.20% | 1 | 3 | Keppra prophylaxis | d60 | |
| PTPN2-KO CAR-T DL3#2 | 4y 8m | Male | 8.5 kg | 1.00E+06 | 27.40% | 2 | 8 | Keppra prophylaxis<br>Tocilizumab x2<br>Dexamethasone x1 | d60 | |
| PTPN2-KO CAR-T DL3#3 | 7y 6m | Female | 8 kg | 1.00E+06 | 5.12% | 1 | 1 | Keppra prophylaxis | d60 | |
| PTPN2-KO CAR-T DL4 | 5y 2m | Female | 8.2 kg | 6.00E+06 | 82.40% | 4 | 17 | Keppra prophylaxis<br>Diazepam x2<br>Tocilizumab x2<br>Dexamethasone x2 | d7 | Severe ICANS |

[1] Refer to Supplemental Table 2 for ICANS scoring criteria

**Supplemental Table 2.** Neurotoxicity score / ICANS for NHP recipients

| General/Behavior | | | | |
|---|---|---|---|---|
| **Comfort/Appearance** | **Food intake** | **Respiratory effort** | **Facial Expression** | **Eye Contact** |
| Normal =0 | Normal =0 | Normal =0 | Normal =0 | Yes = 0 |
| Fair/Quiet = 1 | Decreased = 1 | Decreased rate = 1 | Reacting by closing eyes/grimacing = 1 | Limited = 1 |
| Agitated/Distressed = 2 | None = 2 | Increased rate/effort = 2 | Poor facial expression = 2 | No = 2 |
| | | | Expressionless = 3 | |

| Mental status/Consciousness | | | |
|---|---|---|---|
| **Activity Level** | **Auditory Response** | **Visual Response** | **Eye Movement** |
| Normal =0 | Reacts = 0 | Tracks = 0 | Normal =0 |
| Reduced (uninterested, only with stim) = 1 | Doubtful = 1 | Doubtful = 1 | Intermittent deviation/abnormal movements = 1 |
| Increased (agitated, hyperactive) = 1 | No reaction = 2 | No tracking = 2 | Continuous deviation/abnormal movements = 2 |
| Lethargic = 2 | | | |
| Unresponsive = 3 | | | |

| Motor | | | | |
|---|---|---|---|---|
| **Moves all extremities** | **Focal Weakness** | **Abnormal Movements** | **Tremor** | **Rigors** |
| Yes = 0 | No = 0 | No = 0 | None = 0 | No = 0 |
| No = 1 | Yes = 1 | Yes, suppressible = 1 | Minimal = 1 | Yes = 1 |
| | | Yes, not suppressible = 2 | Mild = 2 | |
| | | | Moderate = 3 | |
| | | | Severe = 4 | |
| **Coordination: On Perch** | **Coordination: On Ground** | **Reaching** | **Gait** | |
| Normal balance = 0 | Normal balance = 0 | Normal (L/R) = 0 | Normal = 0 | |
| Unstable = 1 | Unstable = 1 | Unsteady (L/R) = 1 | Ataxic = 1 | |
| Unable to sit = 2 | Unable to sit = 2 | | | |

| Seizure Activity |
|---|
| No = 0 |
| Yes = 1 |
